# Utilizing random regression models for genomic prediction of a longitudinal trait derived from high-throughput phenotyping

**DOI:** 10.1101/319897

**Authors:** Malachy Campbell, Harkamal Walia, Gota Morota

## Abstract

The accessibility of high-throughput phenotyping platforms in both the greenhouse and field, as well as the relatively low cost of unmanned aerial vehicles, have provided researchers with an effective means to characterize large populations throughout the growing season. These longitudinal phenotypes can provide important insight into plant development and responses to the environment. Despite the growing use of these new phenotyping approaches in plant breeding, the use of genomic prediction models for longitudinal phenotypes is limited in major crop species. The objective of this study is to demonstrate the utility of random regression (RR) models using Legendre polynomials for genomic prediction of shoot growth trajectories in rice (*Oryza sativa*). An estimate of shoot biomass, projected shoot area (PSA), was recored over a period of 20 days for a panel of 357 diverse rice accessions using an image-based greenhouse phenotyping platform. A RR that included a fixed second-order Legendre polynomial, a random second-order Legendre polynomial for the additive genetic effect, a first-order Legendre polynomial for the environmental effect, and heterogeneous residual variances was used to model PSA trajectories. The utility of the RR model over a single time point (TP) approach, where PSA is fit at each time point independently, is shown through four prediction scenarios. In the first scenario, the RR and TP approaches were used to predict PSA for a set of lines lacking phenotypic data. The RR approach showed a 11.6% increase in prediction accuracy over the TP approach. Much of this improvement could be attributed to the greater additive genetic variance captured by the RR approach. The remaining scenarios focused forecasting future phenotypes using a subset of early time points for known lines with phenotypic data, as well new lines lacking phenotypic data. In all cases, PSA could be predicted with high accuracy (*r*: 0.79 to 0.89 and 0.55 to 0.58 for known and unknown lines, respectively). This study provides the first application of RR models for genomic prediction of a longitudinal trait in rice, and demonstrates that RR models can be effectively used to improve the accuracy of genomic prediction for complex traits compared to a TP approach.

## 1 Introduction

With the advent of next-generation sequencing technologies, the biology community has experienced a rapid increase in the amount of genotypic data that is available. These developments, along with the low cost of sequencing, has encouraged the adoption of genomic selection (GS) approaches in plant breeding. With these approaches, genome-wide SNP markers are used to estimate an individuals additive genetic contribution to a given trait, and genotyped individuals can be selected and advanced to further generations without phenotypic evaluation (Meuwissen et al., 2001; Jannink et al., 2010; Endelman, 2011). Although these approaches have increased genetic gain through the acceleration of breeding cycles, considerable resources must still be devoted to the accurate phenotypic evaluation of individuals (Furbank and Tester, 2011). This necessary step remains a major bottleneck for many breeding programs.

In recent years, considerable investment, in both the public and private sector, have been made to automate the phenotypic characterization of large populations. Large investments have been made to build high-throughput phenotyping facilities in both the greenhouse and field where highly controlled water, nutrient, or temperature regimes can be applied to individual plots, and plants can be routinely monitored throughout the development using imaging. Moreover, the relatively low cost of drones that can be fitted with cameras and other sensors, have provided researchers with an effective means to characterize large populations throughout the growing season (Furbank and Tester, 2011; Chapman et al., 2014; Zhang et al., 2016; Watanabe et al., 2017). These longitudinal phenotypes can provide important insight into the mechanisms that underlie physiological responses to environmental stresses and developmental processes, and can be leveraged to improve prediction accuracies for complex polygenic traits, such as yield that have been a target for most breeding programs (Fahlgren et al., 2015; Campbell et al., 2017; Sun et al., 2017). Despite the growing use of these new phenotyping approaches in plant breeding, the use of models for genomic selection (GS) for longitudinal phenotypes is limited in breeding major crop species. Most conventional field studies involve one or a few evaluations throughout the growing season, thus repeated phenotypic measurements on the same plant or plot is relatively rare.

Several approaches have been utilized for GS using longitudinal data. A simple repeatability (SR) model was used by Sun et al. (2017) and Rutkoski et al. (2016) for secondary longitudinal traits. The SR model treats each time point as a repeated measure of the same trait and assumes that the variance for all records are equal and the correlation between time points is constant. However, for many traits recorded across many time points, the assumption behind SR model is not realistic. A multivariate approach can be extended to longitudinal data. However, the computational complexity of the multivariate approach increases with the number of time points, and becomes unfeasible with high frequency longitudinal traits due to the large number of parameters to estimate. Often, the number of observations necessary to accurately estimate parameters exceeds the size of most studies.

Random regression (RR) models have proven to be an attractive alternative to the above methods, and have been utilized in livestock and tree breeding (Apiolaza et al., 2000; Bermejo et al., 2003; Nobre et al., 2003; Bohmanova et al., 2008; Costa et al., 2008; Wetten et al., 2012; Howard et al., 2015). Here, covariance functions are explicitly defined that are equivalent to the full covariance matrix of the trait across time points (Kirkpatrick et al., 1990; Meyer, 1998). Covariance functions include, but are not limited to banded correlation, autoregressive models, orthogonal polynomials, or spline functions (Meyer, 1998; Apiolaza et al., 2000). Thus, these models utilize a few parameters to describe the full covariance, and are much more computationally efficient. In animal breeding, RR models have been used extensively to estimate hertiabilities and perform pedigree-based prediction of important longitudinal traits such as growth, feed intake, fat, and milk production (Bermejo et al., 2003; Nobre et al., 2003; Bohmanova et al., 2008; Costa et al., 2008; Wetten et al., 2012; Howard et al., 2015).

The increased accessibility to high-throughput phenotyping platforms provides the plant science community with high frequency temporal measurements for complex polygenic phenotypes. These data are very different from those typically used for genomic prediction in which phenotypes are recorded at a single time point or at harvest for large populations. However, the availability of these new data presents an opportunity to extend these approaches used extensively for longitudinal traits in animal breeding to major crops. Here, we demonstrate the use of RR models to predict shoot growth trajectories in a rice diversity panel. Specifically, the aims of this study are are to (1) examine the advantage of utilizing longitudinal phenotypes over single end-point measurements (cross-sectional GS), (2) determine whether longitudinal phenotypes collected during early time-points can be used to predict phenotypes at later time points (i.e. forecasting lines with records), and (3) predict future phenotypes for new lines using early records for existing lines.

## 2 Materials and Methods

### 2.1 Plant materials and greenhouse conditions

Three hundred seventy eight lines of the Rice Diversity Panel 1 were selected for this study (Zhao et al., 2011). Seed propagation is described in Campbell et al. (2015). Three uniformly germinated seedlings were selected and transplanted to pots (150mm diameter x 200 mm height) filled with approximately 2.5 kg of UC Mix (the actual weight varied from experiment to experiment by 100-200 g). Square containers were placed below each pot to allow water to collect.

### 2.2 Experimental Design

All experiments were conducted at the Plant Accelerator, Australian Plant Phenomics Facility, at the University of Adelaide, SA, Australia. Each experiment consisted of 378 lines and was repeated three times from February to April 2016. Two smarthouses were used for each experiment, with 216 pots positioned across 24 lanes in each smarthouse. Each experiment consisted of a partially replicated design, with 54 randomly selected lines having two replicates in each experiment.

Seven days after transplant (DAT), plants were thinned to one seedling per pot. Two layers of blue mesh was placed on top of the pots to reduce soil water evaporation. The plants were loaded on the imaging system and were watered to 90% field capacity at 11 DAT.

### 2.3 Image analysis

The plants were imaged daily from 13 to 33 DAT using a visible (red–green–blue camera; Basler Pilot piA2400–12 gc, Ahrensburg, Germany) from two side-view angles separated by 90° and a single top view. The three experiments produced a total of 72,537 images. “Plant pixels” were extracted from RGB images using the LemnaGrid software. Briefly, plant pixels were extracted from background objects using a color classification strategy. Two set of colors were chosen manually to represent plant and background objects. For each image, pixels were assigned as background or plant pixels using the nearest-neighbor method. For a given pixel, this method assigns the pixel to a predefined color by finding the most similar (smallest Euclidean distance) color in the set. Noise (i.e. small areas of non-plant pixels) in the image is removed using a series of erosion and dilation steps.

The sum of the “plant pixels” from the three RGB images were summed, and used as a measure of shoot biomass. Here this trait is referred to as projected shoot area (PSA). This metric has been shown to be an accurate representation of shoot biomass (Campbell et al., 2015; Golzarian et al., 2011; Knecht et al., 2016). Prior to downstream analyses, outlier plants at each time point were detected for each trait using the 1.5(IQR) rule. Plants that were flagged as potential outliers were plotted and inspected visually. Those that exhibited abnormal growth patterns were removed. A total of 32 plants were removed, leaving a total of 2,604 plants for downstream analyses.

### 2.4 Selection of random regression models

PSA was modeled across all twenty time points using several RR models. Following the notation of Mrode (2014), the RR models can be summarized as

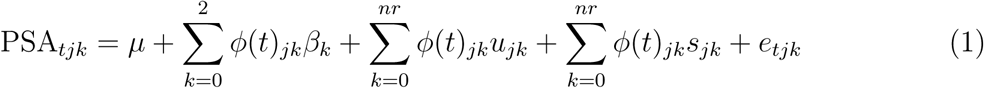

Here *β* is the fixed second-order Legendre polynomial to model the overall trend in the trait overtime, *uj_k_* and *sj_k_* are the *k^th^* random regression coefficients for additive genetic effect and random experiment of line *j*, *nr* is the order of polynomial for the random effects, and *e_tjk_* is the random residual. The order of *β* was selected based on visual inspection of the trends. Various polynomial functions and residual variance structures were evaluated for line and experiment, and residuals, respectively. A complete description of the models is provided in Table 1. For each trait, the models were ranked based on goodness-of-prediction using Akaike’s information criterion (AIC) scores (Akaike, 1974).

### 2.5 Genomic selection at each time point

A mixed model approach was used to fit genomic best linear unbiased predictions (gBLUPs) at each time point using the following model.

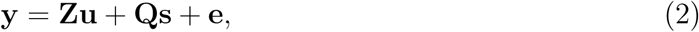

Here, **y** is the PSA at time *t*; **Z** and **Q** are incidence matrices corresponding to the random additive genetic effect (**u**), and random experimental effect (**s**), respectively; and **e** is the random residual error. For the random terms we assume 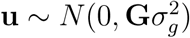, 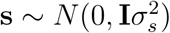, and 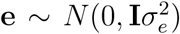. Here, 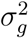 is the additive genetic variance; 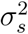 is an environmental variance associated with experiment; and 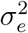 is the residual variance. A genomic relationship matrix (**G**) was calculated using VanRaden (2008).

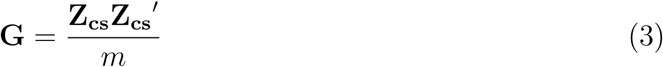

Here, **Z_cs_** is a centered and scaled *n* × *m* matrix, where *m* is 33,674 SNPs and *n* is the 357 genotyped rice lines.

### 2.6 Genomic selection using random regression

For each trait, the “best” random regression model was used to predict gBLUPs. The following mixed model was used to predict gBLUPs

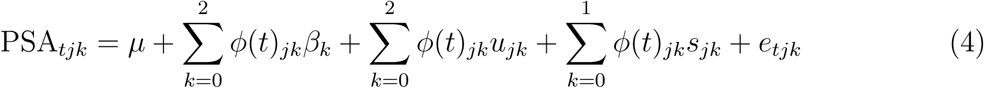

The variables are the same as in *Selection of random regression models*, however note that *nr* has been replaced with 2 and 1 for the additive genetic and experiment effect, respectively. Thus the random additive genetic effects are described using a second-order Legendre polynomial, while a first-order Legendre polynomial is used to describe the experiment effects across time points.

In matrix notation, the model is

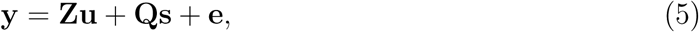
with all vectors and matrices defined as above. However here *u* is now a vector of random regression coefficients for the additive genetic effects. For the random terms we assume **u** ~ *N*(0, **G** ⊗ **Q**), **s** ~ *N*(0,**I** ⊗ **P**), and **e** ~ *N*(0,**I** ⊗ **D**). Here, **Ω** is a 3 × 3 covariance matrix of random regression coefficients for additive genetic effects; **P** is a 2 × 2 covariance matrix of random regression coefficients for experiment effect; and **D** is a diagonal matrix allowing for heterogeneous variances over time points. **Z** and **Q** are covariable matrices where the *i*th row contains the orthogonal polynomials for the *i*th day of imaging. Thus, matrix **Z** is the covariable matrix for the additive genetic effects with a dimension of *t* × *nk* where *nk* is the order of Legendre polynomial for the additive genetic effect multiplied by the number of individuals with phenotypic records and *t* refers to the number of days of imaging. Similarly, **Z** is a *t* × *ns* covariable matrix for the experiment effect, where *ns* is the the order of the Legendre polynomial for the experiment effect (e.g. 1) time the number of experiments (e.g. 3). Variance components and gBLUPs were obtained using ASREML (Release 4.0) (Gilmour et al., 2015).

Using the method above, variance components were obtained for additive genetic and environmental components. For the additive genetic term, each line has three random regression coefficients (*nr* = 0,1, 2). gBLUPs were predicted at each time point according to Mrode (2014). For a given line, *j*, at time *t* the gBLUPs can be obtained by 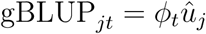; where *ϕ_t_* is the row vector of the matrix of Legendre polynomials of order 2.

### 2.7 Estimation of narrow-sense heritability

To estimate the narrow sense heritability, variance components were obtained for each random term using ASREML for the TP analyses and the RR approach. For the RR approach, additive genetic variance was obtained at each time points using methods described by Mrode (2014). Briefly, for time *i* the genetic variance can be obtained by 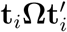, where **t***_i_* = *ϕ_ik_*, the *i*th row vector of the matrix of Legendre polynomials at different time points (*ϕ*) for the *i*th day of imaging, **Ω** is the covariance matrix of RR coefficients for the genetic effects, and *k* is the order of fit. The variance of the experimental effect across time points was calculated using the same approach. For both the single time point analysis *h*^2^ was estimated as 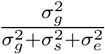.

### 2.8 GS scenarios and cross validation

Four scenarios were tested using GS (Figure 1). In the first scenario (scenario A), all twenty time points were used to fit a RR model and phenotypes were predicted for a set of lines without phenotypic records. The second scenario (scenario B), the dataset was split into two datasets each consisting of ten consecutive time points. A RR model was fitted using the first ten time points and was used to predict the phenotypes for the same set of lines in the last ten time points. Scenario C, can be thought of as a combination of scenarios A and B. Here, the dataset was split into four subsets, with each quadrant consisting of 178 to 179 lines and ten time points. Here, a RR model was fitted using ten early time points for half the lines with known phenotypes, and was used to predict the phenotypes in the last ten time points for the remaining 178 to 179 lines. Finally, in the last scenario (Scenario D) we sought to predict the shoot biomass at a later time points in an independent study. This can be thought of as forecasting for new lines in an independent study. A publicly available dataset was used in which 359 lines (357 lines in common between the two studies) were phenotyped from 20 to 40 days after transplant, thus a 13 day overlap was available for the two datasets, and a RR model was fitted using phenotypic information from the time points in the first experiment for 179 lines, and was used to predict gBLUPs for the remaining 178 lines in a second independent experiment described by Campbell et al. (2017).

**Figure 1:**
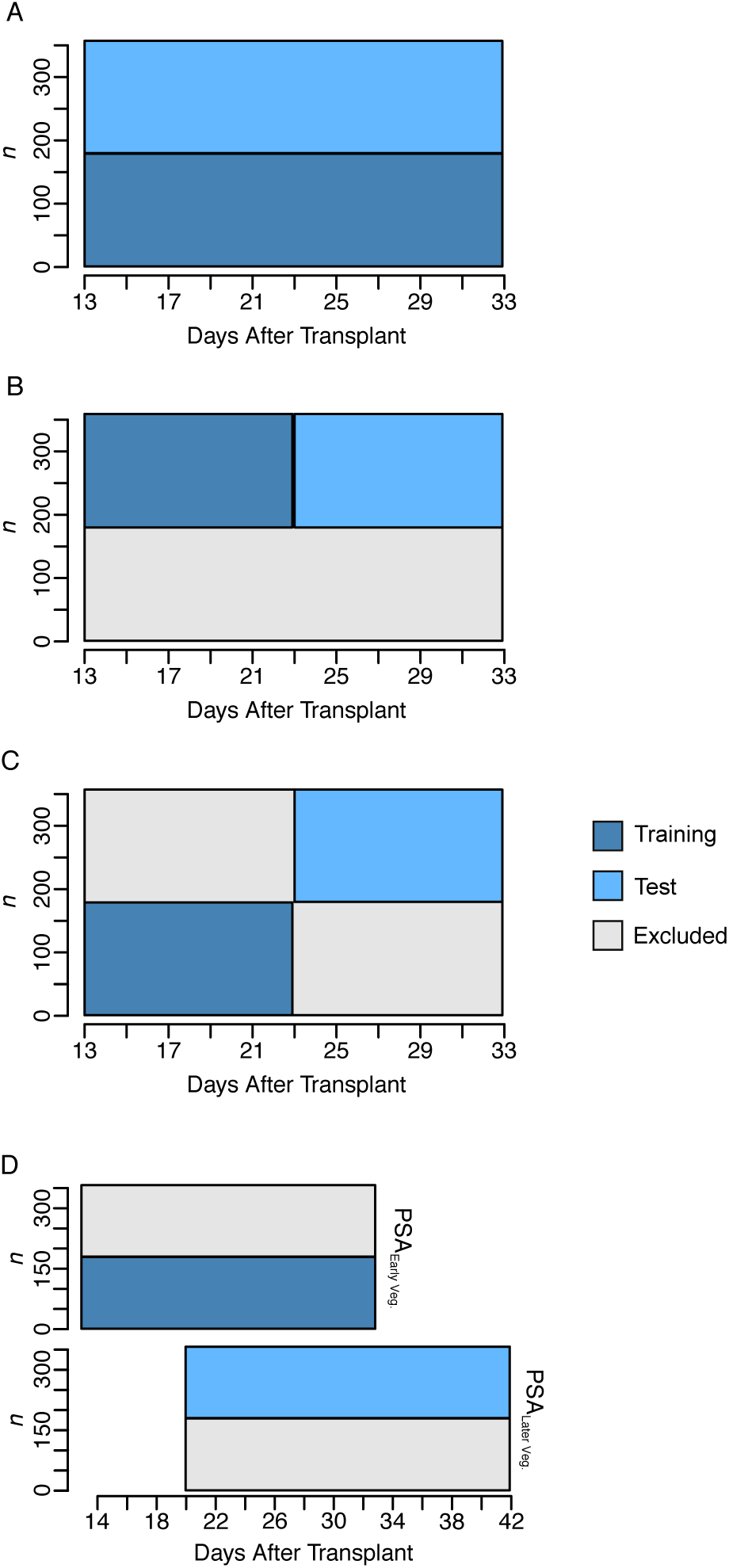
Graphic representation of cross validation schemes for predicting longitudinal phenotypes using random regression. In (A), (C), and (D) two-fold cross validation was used, where phenotypes for 179 lines were used as a training set to predict phenotypes for the remaining 178 lines. In (A), all twenty time points for the training set were used to predict the phenotypes at each of the twenty time points for an new set of lines. The second scenario (B) can be thought of as a forecasting approach where the dataset was split into two longitudinal datasets each consisting of ten time points. The first ten time points for 179 lines and were used to predict the phenotypes at the last ten time points for the same 179 lines. In (C), a forecasting approach was again used, however the lines were randomly split in two, and the first ten time points were used to predict phenotypes in the last ten time points for a group of new lines. In (D) the first 20 time points was used to predict gBLUPs at a later time points in an independent study. Here, a publicly available dataset was used as a testing set in which 357 lines were phenotyped from 20 to 40 days after transplant, thus a 13 day overlap was available for the two datasets. Here, the independent dataset is indicated with *PSA_LaterVeg._*. Excluded indicates that these data points were not included for analyses.

To assess the accuracy of gBLUPs for the TP GS as well as scenarios A, C, and D, a two-fold cross validation approach was used. Briefly, the 357 lines were split into two sets, with one serving as a training set with known phenotypes and the second serving as a testing set with unknown phenotypes. Since the number of lines were not even the remaining line was assigned to the training set. The accuracy of prediction was assessed by comparing predicted gBLUPs with observed PSA at each of the three experiments using Pearson’s correlation method. The lines were randomly assigned to each fold, and the process was repeated 20 times. For each fold, the average correlation over the three experiments was used, and the average over the two folds was used for each resampling run. For scenario B, half of the lines were randomly selected and the first ten time points were used to predict the phenotypes in the last ten time points for the same lines. Again, the variance in prediction accuracy was assessed by randomly sampling half the lines for analysis. Pearson’s correlation was computed for the gBLUPs and PSA as described above.

## 3 Results

A rice diversity panel was phenotyped over a period of twenty days during the early vegetative stage using an automated high-throughput phenotyping platform. The panel consists of 357 lines from 80 countries, and captures much of the genetic diversity within cultivated rice (Zhao et al., 2011).

The plants were imaged each day using RGB cameras from three angles (two side view angles separated by 90 degrees and one top view). The plant pixels from each image were summed and used to estimate shoot biomass. Here, this metric is referred to as PSA and has been shown to be an accurate measure of shoot biomass in cereals (Berger et al., 2010; Campbell et al., 2015). This experiments captures the early vegetative stage of development, where shoot biomass increases nearly exponentially (Figure 2A, Figure S1).

**Figure 2:**
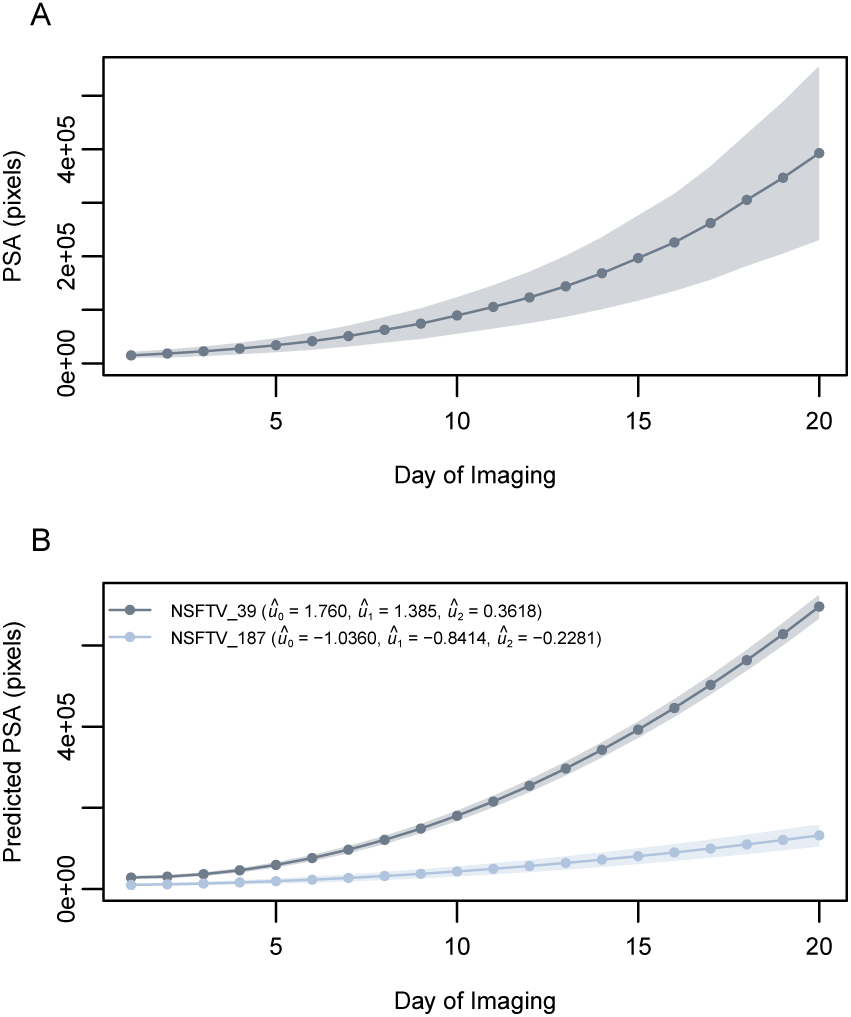
Projected shoot area (PSA) across twenty days of imaging. (A) Population mean for PSA across the twenty days of imaging. Here, the shaded region represents the standard deviation of PSA at each time point. (B) Predicted PSA for two contrasting lines using a random regression (RR) model. The RR model included a fixed second-order polynomial to model the mean trend in shoot growth, a second-order Legendre polynomial for the random additive genetic effect, a first-order Legendre polynomial for the experimental effect, and the residual variance was assumed to be heterogeneous over time points. The predicted RR coefficients for each line are provided in the figure legend. The shaded regions represent the standard error of predicted PSA at each time point. Here, PSA is defined as the sum of plant pixel from three images (two side-view images and one top-view). The shaded region represents the standard deviation of PSA at each time point.

### 3.1 Random regression model selection

RR models have been used extensively to model longitudinal phenotypes in animal breeding. These models are particularly advantageous in that differences in the shape of the curve can be accounted for, and can be solved using the conventional mixed model framework. Thus, in the scope of genetics, these models allow for inter-individual variation in the mean trend to be estimated. Here, the overall mean growth trend was modeled using a second-order Legendre polynomial. A total of eight models were evaluated to identify a model that adequately described the data and could be used for GS. Each model included a fixed second-order Legendre polynomial to describe the overall mean growth trend, while several Legendre polynomials ranging from zero to second-order Legendre polynomials were fitted for random genetic and experimental effects. The residual effects were assumed to be constant or heterogeneous across time points using an identity or diagonal matrix, respectively. The “best” model was selected based on the smallest AIC value. Table S1 provides an overall summary of the models and the corresponding AIC values. The “best” model (Model 8) was one that included a fixed second-order polynomial to model the mean trend in shoot growth, a second-order Legendre polynomial for the random additive genetic effect, a first-order Legendre polynomial for the experimental effect, and the residual variance was assumed to be heterogeneous over time points. Figure 2B shows the predicted PSA obtained with model 8 for two lines with contrasting contrasting genetic values for the RR coefficients.

### 3.2 Genetic correlation and narrow sense heritability of PSA

To examine the relationship for PSA between time points, the phenotypic and genetic correlation was estimated. Estimates for the overall phenotypic correlations were high (*r*: 0.49 - 1.0), with the highest correlation observed between adjacent time points (Figure 3A). The genetic correlation followed a similar patten, with an overall high correlation (*r*: 0.84 - 1.0) observed among pairwise comparisons of all 20 time points. As above, adjacent time points exhibited the highest genetic correlation (*r* = 1), while those further apart exhibited lower correlation (Figure 3B). Interestingly, a strong genetic correlation was observed between day 1 and day 20 (*r* = 0.91), indicating that shoot growth (e.g. PSA) may be driven by similar genetic mechanisms at the early seedling and active tillering stage in rice.

**Figure 3:**
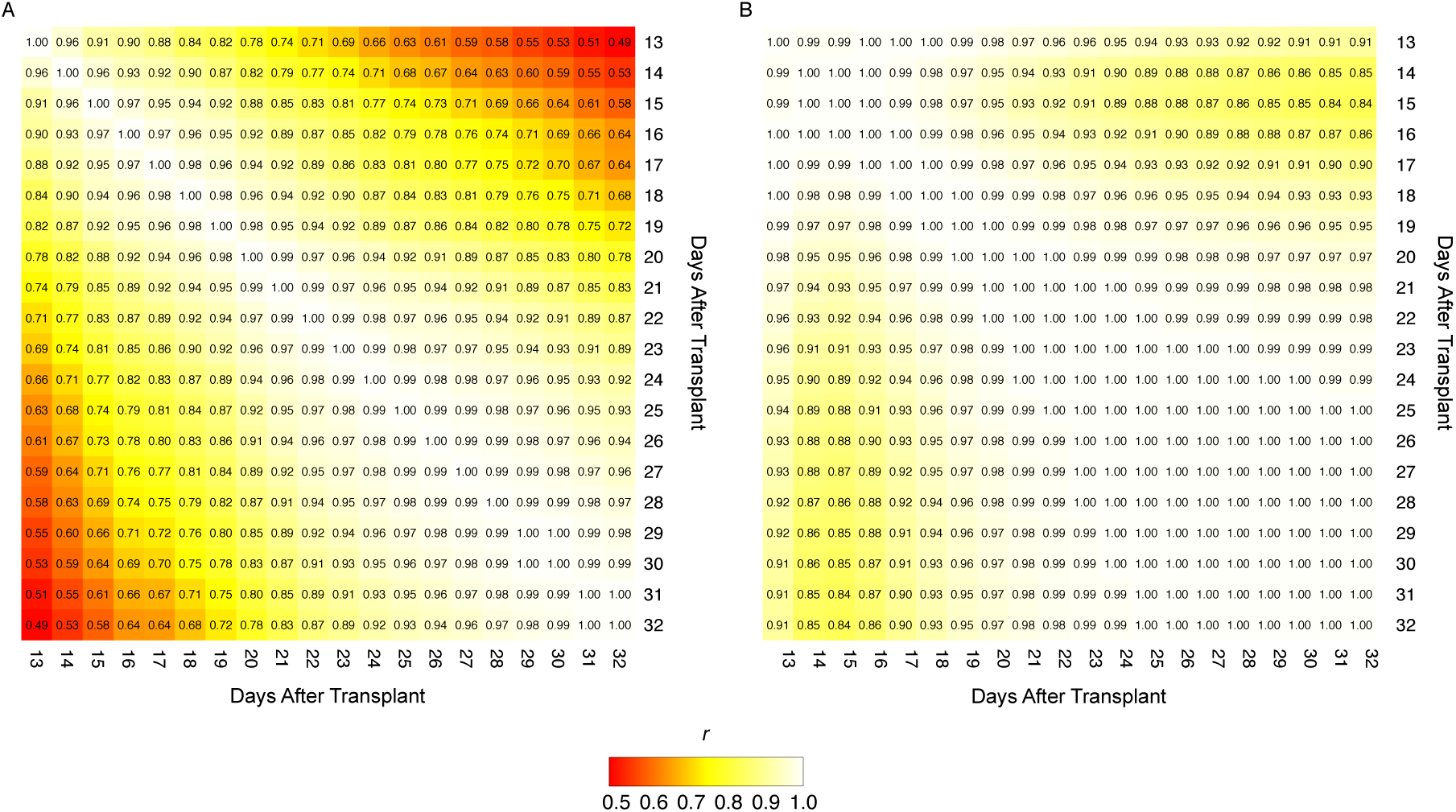
Phenotypic and genetic correlations between each time point. (A) Phenotypic correlations were estimated between time points using Pearson’s method. (B) The inferred genetic correlation matrix of random regression terms for the additive genetic effects were used to estimate the genetic correlations between time points. The scale on the left of each panel indicates the strength of the correlations (*r*).

To evaluate the ability of the longitudinal RR approach to capture additive genetic variance, the narrow sense heritability of PSA was estimated using the RR model described above and a conventional mixed model at each time point. The mixed model included random terms for the additive genetic and experimental effect. For both models, a genomic relationship matrix was generated using 33,674 markers for the 357 lines. On average, the RR approach showed a 44% increase in the heritability of PSA compared to the TP approach (Figure 4). The TP approach showed a mean *h*^2^ of 0.50 over all time points, while the RR approach showed an *h*^2^ of 0.71 on average. *h*^2^ ranged from 0.60 to 0.77 for the RR approach, while *h*^2^ ranged from 0.46 to 0.57 for the TP approach. The two approaches showed nearly identical *h*^2^ estimates on day 1, however at later time points *h*^2^ of RR was considerably higher than TP. These results suggest that the RR approach captures more additive genetic variance for PSA than the TP approach.

**Figure 4:**
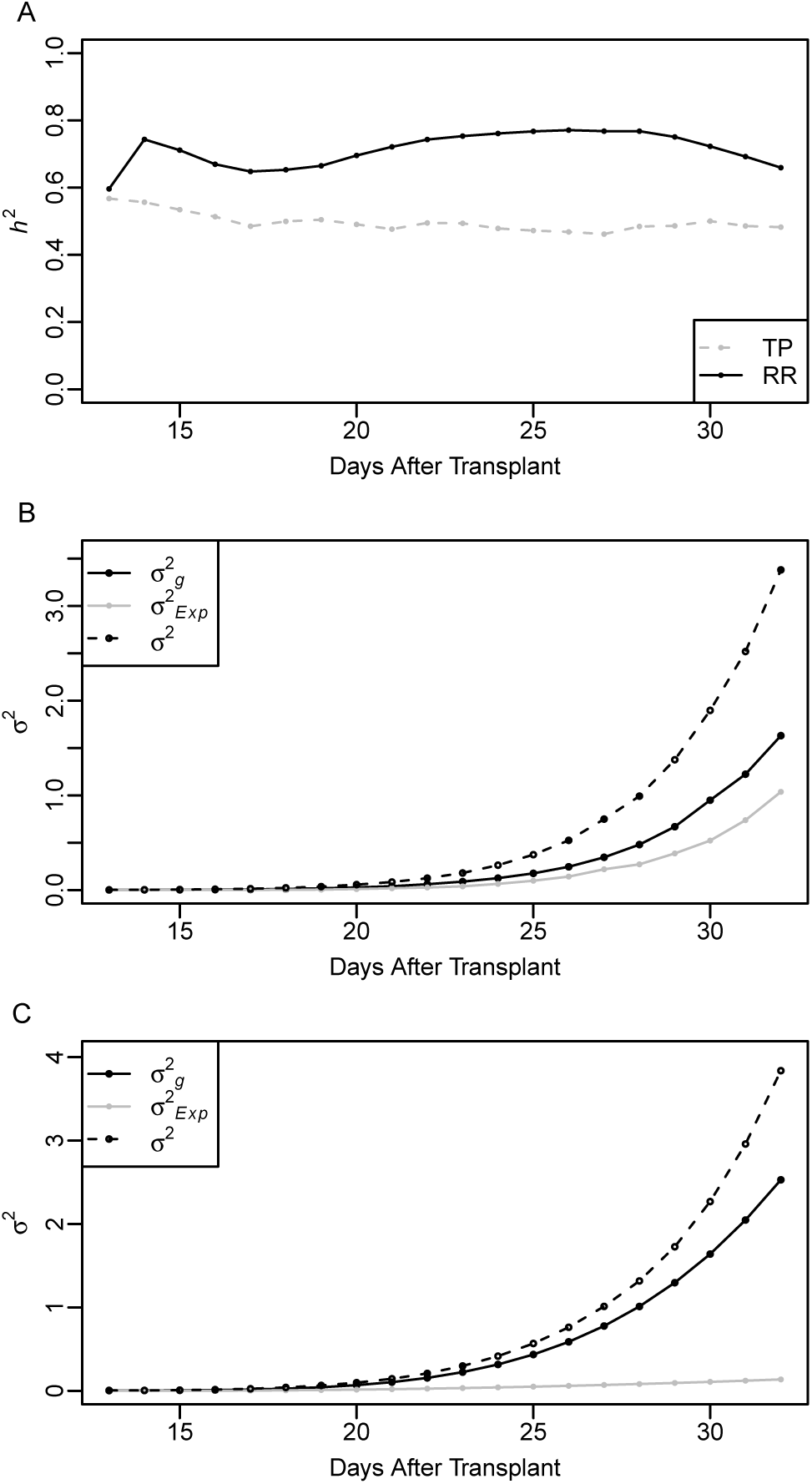
Narrow sense heritability and variance components estimated using the single time point (TP) and random regression (RR) approaches. The narrow sense heritability (*h*^2^) is presented in panel A. Variance components for the TP and RR approaches are pictured in panels B and C, respectively. For the single time point analysis, a conventional mixed model was used to estimate the narrow sense heritability of PSA at each time point. The TP model included a random additive genetic effect and experimental effect. The RR model included a fixed second-order Legendre polynomial, the random additive genetic effect were modeled using a second-order Legendre polynomial, a first-order random effect was used for experiment, and the residual variance was assumed to be heterogeneous over time points. For both models, the experimental term was considered as an environmental effect.

### 3.3 Utility of longitudinal phenotypes for genomic prediction

The availability of high throughput phenotyping platforms provides a means to accurately phenotype large populations for a number of traits throughout time. While phenotypes recorded at a high frequency over time will likely improve the accuracy of GS, few reports have demonstrated the advantages of longitudinal phenotypes in major crops or model plant systems. Here, the utility of longitudinal phenotypes for GS was evaluated under four hypothetical scenarios (Figure 1). The first scenario can be thought of as a standard GS approach (Figure 1A). Here, all 20 time points for half of the 357 lines used to predict the phenotypes at all 20 time points for the remaining lines. The aim of scenario A is to determine whether the longitudinal RR approach provides greater prediction accuracy than a cross-sectional GS approach in which a mixed model is fit at each time point. The first training set can be thought of as existing lines with phenotypic records and the test population as a new set of lines without records. The aim of scenario B (Figure 1B), is to determine if traits at later time points can be predicted for known lines using information at early time points. Thus, it can be considered as a forecasting approach. Here, longitudinal phenotypes are available for lines during the early time points (1-10 days of imaging), and are used to predict phenotypes for the same lines at later time points. Scenario C (Figure 1C), can also be considered a forecasting approach however for new lines. Here a subset of lines with phenotypes during the first 10 time points are used to predict the phenotypes for new lines without phenotypes at the later time points. In scenario D (Figure 1D), we sought to predict the shoot biomass at a later time points in an independent study. Here, a publicly available dataset was used in which 359 lines (357 lines in common between the two studies) were phenotyped from 20 to 40 days after transplant, thus a 13 day overlap was available for the two datasets. A RR model was fitted using phenotypic information from the time points in the first experiment for 179 lines, and was used to predict gBLUPs for the remaining 178 lines in the second experiment.

#### Scenario A: Comparison between longitudinal RR and cross-sectional GS

To evaluate the advantages of using the longitudinal phenotype for PSA for GS over a single time points, the prediction accuracy of the RR model described above was compared to a conventional cross-sectional approach in which the additive genetic effects were estimated at each time point. For both approaches, two-fold cross validation was performed in which half the lines were randomly selected as a training set, and the remaining half was used for prediction. Pearson’s correlation was used to assess the accuracy between predicted gBLUPs and observed PSA in the test set for each experiment. The average correlation across all three experiments was determined for each fold. The resampling process was repeated ten times.

Overall, the RR model showed significantly higher predication accuracies than the TP approach (Figure 5A). On average, the longitudinal phenotype improved prediction accuracy by 11.6% (mean across all time points) compared to the TP approach. The prediction accuracies for the TP approach ranged from 0.40 to 0.60, while for the RR approach accuracies ranged from 0.47 to 0.58. Although the TP approach exhibited low prediction accuracies during the early time points and increasing prediction accuracies toward the end of the study, the prediction accuracy for the RR model remained relatively constant with a slight increase in r observed from day 1 to 9. The largest improvements in prediction accuracy was observed between 5 to 10 days of imaging, with the RR model showing 35% higher accuracy at day 8 compared to the TP approach. Collectively, these results indicate that RR models can be used to improve the accuracy of genomic prediction for longitudinal phenotypes.

**Figure 5:**
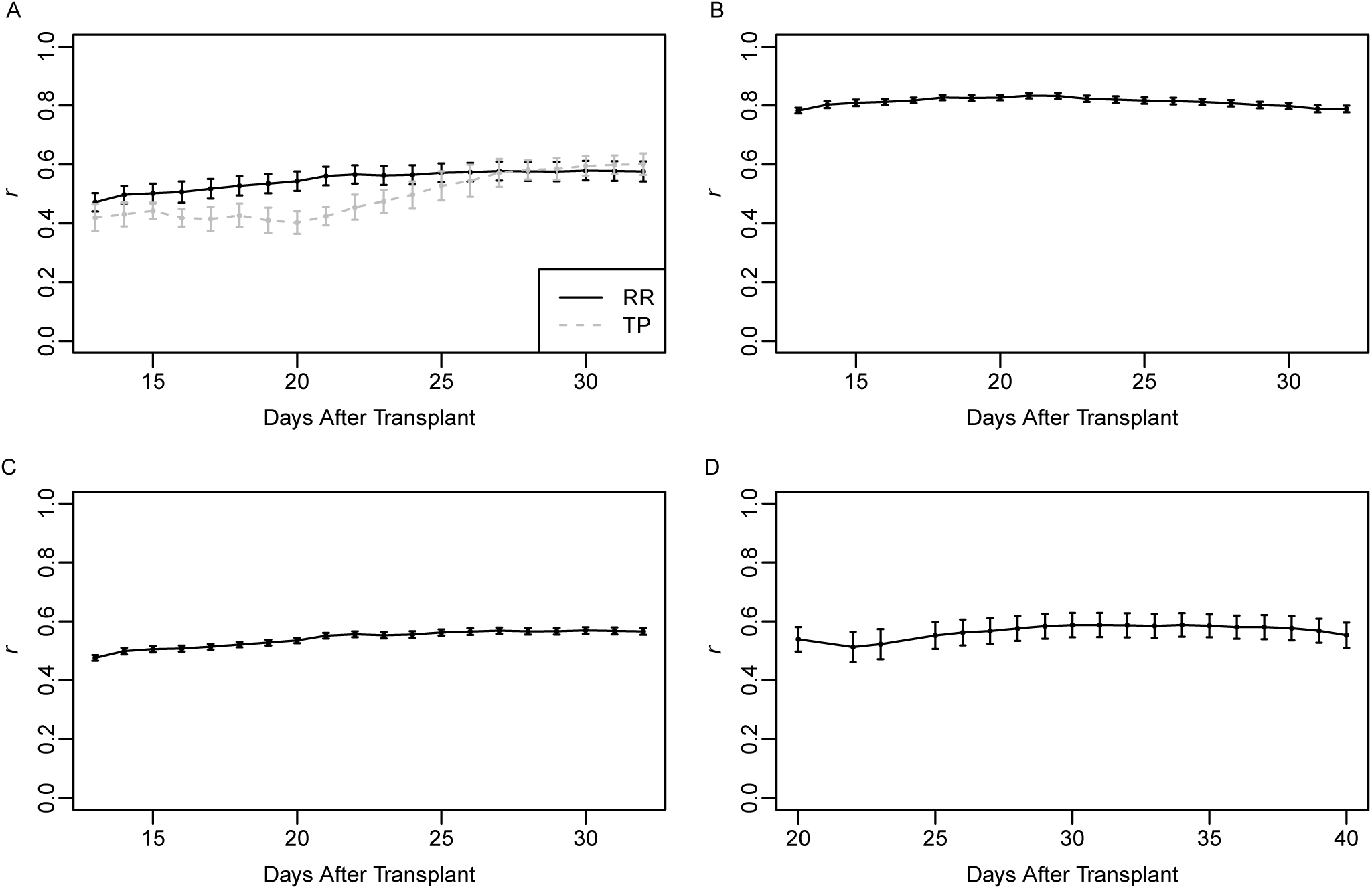
Prediction accuracies of scenarios A to D. For the random regression (RR) approach, a RR model was fit using phenotypic records for 178-179 lines over 20 days. A univariate single time point (TP) run using phenotypic records for 178-179 lines at each day. In both cases, genetic effects from each model were used to predict gBLUPs for the remaining 178-179 lines. Prediction accuracy was assessed using Pearson’s correlation between the predicted gBLUPs and observed PSA for the test set. Resampling was done twenty times. The error bars represent the standard deviation where *n* = 20. A comparison of prediction accuracies for TP and RR approaches is presented in (A). Panels B and C present the prediction accuracies for forecasting future phenotypes using phenotypic information at early time points for known lines (B) and new lines (C). Panel D provides prediction accuracies for forecasting future phenotypes in an independent study using phenotypes from an earlier developmental period.

#### Scenario B: Forecasting existing lines

Here, the the objective is to predict future phenotypes for lines with phenotypic trajectories recorded earlier in the growing season or development. To this end, the dataset was separated into two, with the first ten time points serving as a training set to predict the phenotypes for the last ten days. This approach is described in Figure 1B. The RR model described above was fit to the data. To assess the accuracy of prediction, two-fold cross validation was performed in which 50% of the lines were randomly selected for training and prediction, and the resampling process was repeated ten times. The accuracy of prediction was very high, ranging from 0.79 to 0.82 for the last ten time points without phenotypic records (Figure 5B). A slight decline in prediction accuracy was observed after day 10, with day 11 exhibiting the highest accuracy (*r* = 0.82) and the lowest accuracy on day 20 (*r* = 0.79). This trend in prediction accuracy is expected, given that the phenotypic records at day 11 should be very highly correlated with those at day 10, with the correlation declining as time progresses. The high predictive ability observed indicates that the first ten time points is sufficient to accurately predict future phenotypes for known lines.

#### Scenario C: Forecasting new lines

As shown above, future phenotypes can be accurately predicted from longitudinal traits at early time points for existing lines. While the knowledge of performance of known lines at future time points may be beneficial in some applications, GS is most often used to select lines without prior knowledge of the phenotype. Previously in scenario A, we showed that phenotypes could be predicted accurately for new lines using the complete longitudinal phenotype. Here, the aim is to predict future phenotypes for new lines with no phenotypic records using early phenotypic records for existing lines. To this end, the dataset was partitioned into two temporal datasets, with the first ten time points serving as a training set to predict the phenotypes for the last ten days (Figure 1C). As above, a two-fold cross validation approach was used to assess prediction accuracy. Half the lines were randomly assigned to each fold, and the first ten time points from the first fold were used to predict the phenotypes at the last ten time points in the second fold. The prediction accuracies for scenario C were very similar to those observed for scenario A. Accuracies ranged from 0.48 to 0.57, with the prediction accuracy ranging from 0.55 to 0.57 in the last ten days (Figure 5C). The prediction accuracies showed a slight increase from day 1 to day 9. The highest prediction accuracy was observed at day 15, while the lowest accuracy was observed at day 1. These results suggest that future phenotypes can be forecast for new lines with reasonable accuracy using phenotypic records from earlier time points for a set of known lines.

#### Scenario D: Forecasting new lines at later time points in an independent study

In scenario C, we have shown that gBLUPs for new lines can be accurately predicted using phenotypes for a set of known lines at a subset of early time points. Here, the objective was to expand this approach and evaluate the utility of the RR model to predict gBLUPs for new lines at future time points in an independent study. Here, we utilized an existing dataset where 359 lines from the Rice Diversity Panel 1 were phenotyped from 20 to 40 days after transplant (Figure 1D.). Although there is overlap between developmental stages of this dataset and the dataset used for scenarios A-C, this experiment was conducted at a different time of year and therefore the photoperiod and light intensity should be different between the two.

A RR model was fitted that was identical to that used for scenarios A-C, in that it included a fixed second-order polynomial to model the mean trend in PSA, a second-order Legendre polynomial for the random additive genetic effect, a first-order Legendre polynomial for the random experimental effect, and a heterogeneous residual variance over time points. The RR model was fitted using phenotypes for 179 lines from the early vegetative stage experiment (i.e. 13 to 32 DAT), and the genetic values for the RR coefficients were used to predict the phenotypes for the remaining 178 lines in the second experiment (i.e. 20 to 40 DAT). A two-fold cross validation approach was used in which phenotypes across all twenty days were selected for 179 lines in the first experiment and were used to predict gBLUPs for the remaining 178 lines in the second experiment.

The prediction accuracy was high with r values ranging from 0.51 to 0.59 (Figure 5D). The prediction accuracy was relatively constant, but showed a slight increase in accuracy from 22 to 29 days after transplant. An increase in the prediction accuracy was observed from 13 to 31 DAT, after which the prediction accuracy declined slightly. The second time point (22 DAT) exhibited the lowest prediction accuracy (*r* = 0.51). The highest prediction accuracy was observed on day 34 after transplant (*r* = 0.59). Collectively, these results suggest that longitudinal phenotypes can be accurately predicted in an independent study using the RR approach.

### 3.4 Discussion

High-throughput phenotyping platforms provide an accessible means to record traits non-destructively for large populations throughout development. Such longitudinal data provide an opportunity to understand the genetics of the development of a phenotype, and identify individuals that exhibit desirable trait trajectories. However, such data provides new challenges to adapt approaches utilized for single time point phenotypes in plant genomics and breeding to accommodate longitudinal data. This study provides the first application of RR models for genomic prediction of a longitudinal trait in rice.

#### Advantages of RR over univariate genomic prediction

The predictive ability in GS is dependent on the heritability of the trait, the number of markers, population size, linkage disequilibrium (LD), and the number of QTL influencing the trait (Daetwyler et al., 2008, 2010). Here, the RR model using longitudinal phenotypes provided greater prediction accuracy compared to the TP gBLUP. The predictive ability of the RR approach improved prediction accuracies by 11.6% on average compared to TP analysis. The number of markers, population size, LD, and the number of QTL influencing PSA are held constant between the two models. Thus, the difference in prediction accuracy hold be largely attributed to the differences in heritability between the RR approach and TP analysis. As shown in Figure 4, the RR approach accounted for more additive genetic variance than the TP analysis. Similar gains in heritability for height in Swedish Scots pine has been reported by Wang et al. (2009) with RR models that utilize B-splines or Legendre polynomials over TP analyses. Moreover, when the prediction accuracy is expressed as the ratio of the correlation of gBLUPs and observed PSA to the square root of *h*^2^, both approaches were nearly equivalent (Figure S2). Thus, the higher prediction accuracy is due to the higher *h*^2^ of the RR approach relative to the TP approach.

With both methods (RR and TP), we observed high prediction accuracies ranging from 0.4 - 0.6 (Arruda et al., 2015; Duhnen et al., 2017; Kristensen et al., 2018; Leplat et al., 2016). While similar accuracies have been reported by other studies for complex traits, it is important to note that the current study utilized a diversity panel with considerable population stratification and the prediction models did not account for population structure. Accounting for population structure is important in genome wide association studies to reduce spurious associations (Yu et al., 2006). However, these corrections can often hinder the ability to detect true QTL that are correlated with population structure (Zhao et al., 2011). With GS, the aim is to achieve high prediction accuracies across subpopulations rather than to detect QTL associated with the trait (Hayes et al., 2009; Lorenz et al., 2011). Thus, the high prediction accuracies observed for the models used in this study may be due, in part, to population structure, however the random sampling of individuals across subpopulations during CV should reduce the possibility of having a training set that is strongly imbalanced by a given subpopulation.

#### Utilizing RR prediction for forecasting phenotypes

The utilization of genomic information to predict future outcomes is not new. Considerable effort in the field of personalized medicine has been devoted to predict disease risks for individuals based on genomic information. Here, disease-associated loci are used to predict potential future outcomes for individuals (Moser et al., 2015). The ability to predict future phenotypes using phenotypic information collected early in the life cycle may be advantageous in plants, particularly perennial species with long life cycles. Selection during the early developmental stage can shorten evaluation times.

Here, we evaluated the ability of RR models to predict future phenotypes using phenotypic records collected during the early time points. This was performed for known lines (e.g. those with early records; Scenario B), as well as new lines (Scenario C and D). We observed high prediction accuracies for each forecasting scenario. As expected the highest accuracy was observed for Scenario B, in which early phenotypic records are used to predict future phenotypes for the same set of lines. Surprisingly, high prediction accuracies were also observed when early records for known lines were used to predict future phenotypes for unknown lines (Scenarios C and D). In both cases, the accuracies were not significantly different from those achieved when using phenotypic information for all time points. These results collectively indicate that the future phenotypes can be accurately predicted using a subset of the temporal phenotypes. While these results are encouraging, these forecasting approaches will be highly dependent on the temporal genetic architecture of the trait. The lack of decline by utilizing only a subset of time points is likely due to the high genetic correlation observed between time points. The similar genetic architecture between the early and late time points that is evidenced by the strong positive genetic correlation (Figure 3B) estimated between early (1-10 days) and late (11-20 days) time points. Thus, we suggest to first evaluate the genetic correlation between time points for the trait of interest before utilizing such forecasting approaches.

### 3.5 Conclusion

High throughput phenomics platforms have provided the plant science community with a means to generate high resolution temporal phenotypes for large populations at a relatively low cost. RR models that utilize Legendre polynomials provide a flexible for genomic prediction of longitudinal traits. These approaches provide several advantages over single time point analyses: (1) these models account for more additive genetic variance compared to the TP analysis, which translates to higher predictive accuracies; (2) future phenotypes can be accurately predicted using phenotypic information for earlier time points for known and unknown lines. TP

---

**Figure S1:**
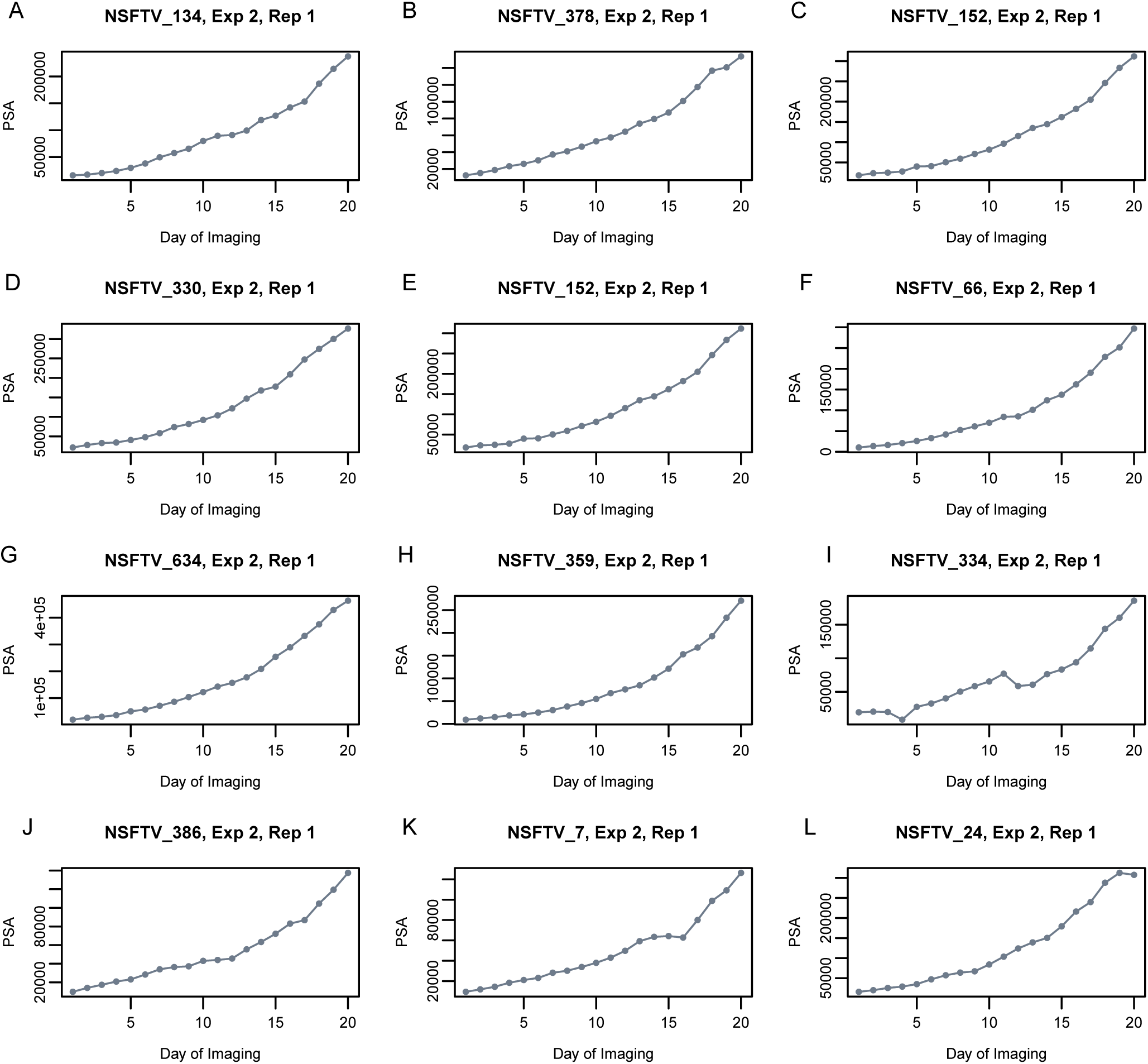
Projected shoot area for a subset of 12 lines. The line identifier (NSFTV_), experiment (Exp), and replicate (Rep) are provided in the plot titles.

**Figure S2:**
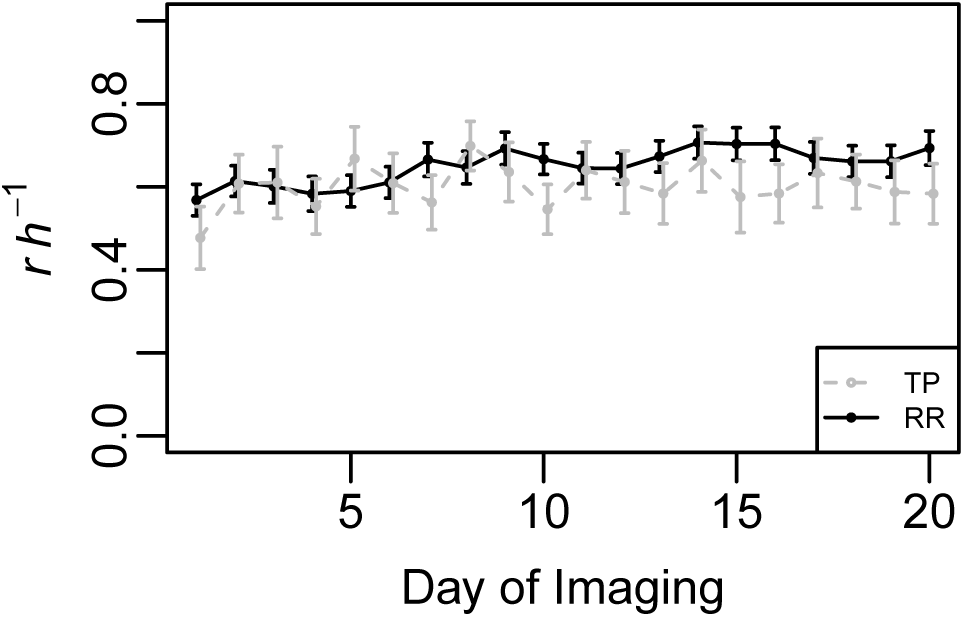
Predictive ability of the random regression (RR) and single time point (TP) approaches expressed as a function of heritability: The analysis followed the same approach as that for scenario A, however for each fold the correlation between gBLUP and observed PSA was divided by the square root of heritability. The error bars represent the standard deviation where *n* = 20.

## Acknowledgements

Funding for this research was provided by the National Science Foundation (United States) through Award No. 1238125 to Harkamal Walia, and Award No. 1736192 to Harkamal Walia and Gota Morota.

## Data Accessibility

The full datasets and all code used in this study is available via GitHub (https:github.com/malachycampbrandom-regression-models-for-genomic-prediction-of-a-longitudinal-trait-derived-from-HTP) and the WRCHR website (WRCHR.org).

## Supplemental Data

**Table S1:** Random regression model selection. Each of the four random regression models included a fixed second-order polynomial to model the mean trend in PSA over the twenty time points, indicated by the column *f*. *G* refers to the random additive genetic effect, *Exp* the random experimental effect, and *e* error term. Models with *Diag* assumed heterogeneous residual variance over time points, while those with *I* assumed the residual variance was constant. *pol^n^* refers to a Legendre polynomial of order *n*.

| *Model* | *f* | *G* | *Exp* | *e* | *LogREML* | *AIC* | *BIC* |
| --- | --- | --- | --- | --- | --- | --- | --- |
| Model 1 | $pol^2$ | $pol^0$ | $pol^0$ | $I$ | 2026.97 | -4047.93 | -4023.65 |
| Model 2 | $pol^2$ | $pol^0$ | $pol^0$ | $Diag$ | 19358.83 | -38673.65 | -38495.60 |
| Model 3 | $pol^2$ | $pol^1$ | $pol^0$ | $I$ | 7345.85 | -14681.69 | -14641.23 |
| Model 4 | $pol^2$ | $pol^1$ | $pol^0$ | $Diag$ | 23273.62 | -46499.24 | -46305.01 |
| Model 5 | $pol^2$ | $pol^2$ | $pol^0$ | $I$ | 8204.64 | -16393.28 | -16328.54 |
| Model 6 | $pol^2$ | $pol^2$ | $pol^0$ | $Diag$ | 24718.93 | -49383.86 | -49165.35 |
| Model 7 | $pol^2$ | $pol^2$ | $pol^0$ | $I$ | 12700.64 | -25381.28 | -25300.35 |
| Model 8 | $pol^2$ | $pol^2$ | $pol^1$ | $Diag$ | 27537.59 | -55017.19 | -54782.49 |

